# Sleep spindle refractoriness segregates periods of memory reactivation

**DOI:** 10.1101/235606

**Authors:** James W. Antony, Luis Piloto, Margaret Wang, Paula Pacheco, Kenneth A. Norman, Ken A. Paller

## Abstract

The stability of long-term memories is enhanced by reactivation during sleep. Correlative evidence has linked memory reactivation with thalamocortical sleep spindles, although their functional role is poorly understood. Our initial study replicated this correlation but also demonstrated a novel rhythmicity to spindles, such that spindles are less likely to occur immediately following other spindles. We leveraged this rhythmicity to test the role of spindles in memory by using real-time spindle tracking to present cues inside versus outside the presumptive refractory period; as predicted, cues presented outside the refractory period led to better memory. Our findings reveal a previously undescribed neural mechanism whereby spindles segment sleep into two distinct substates: prime opportunities for reactivation and gaps that segregate reactivation events.

**One Sentence Summary:** The characteristic timing of sleep spindles regulates when memories can be reactivated during sleep.

---

Memories of daytime episodes are covertly reactivated during sleep, improving memory storage in the brain (*1*). Previous research has implicated three electrophysiological signals in memory processing during sleep. The slowest of these are sleep slow oscillations (SOs), brain rhythms at approximately 1 Hz prominent during deep sleep (*2*). A second signal is the sleep spindle, a burst of activity at 11-16 Hz lasting 0.5-3 s. Third, replay of newly formed memories is thought to occur in conjunction with high-frequency bursts of hippocampal and cortical activity called ripples (*1, 3, 4*). These three signals can occur with precise temporal interrelationships; spindles tend to occur most often during the up-state phase of SOs, and ripples tend to occur at spindle troughs (*5–7*). To the extent that these relationships are evidenced, memory consolidation appears to be more effective (*8*), suggesting a dual cross-frequency coupling mechanism by which initially hippocampal-dependent memories become stabilized in long-term neocortical networks over time (*2*). Pharmacological evidence suggests spindles promote memory consolidation in humans (*9*); however, the time course relating spindles to memory consolidation has not been well-characterized.

Here, we investigated and manipulated temporal relationships between spindles and learning-related auditory cues known to boost memory, relying on a technique called targeted memory reactivation (TMR) (*10, 11*). In experiment 1 (*N* = 18; Fig 1A), subjects first over-learned novel associations between individual sounds and picture items (e.g., [meow]-Brad Pitt, [violin]-Eiffel Tower). Next, they learned unique locations for each item on a background grid. After an initial test, they took an afternoon nap with background white noise (~40 dB; Table S1 provides sleep stage information). Upon online indications of slow-wave sleep (SWS), we embedded half of the cues in the noise, one every 4.5 s. After a post-nap break, subjects took tests on both item-location associations and sound-item associations.

**Figure 1.**
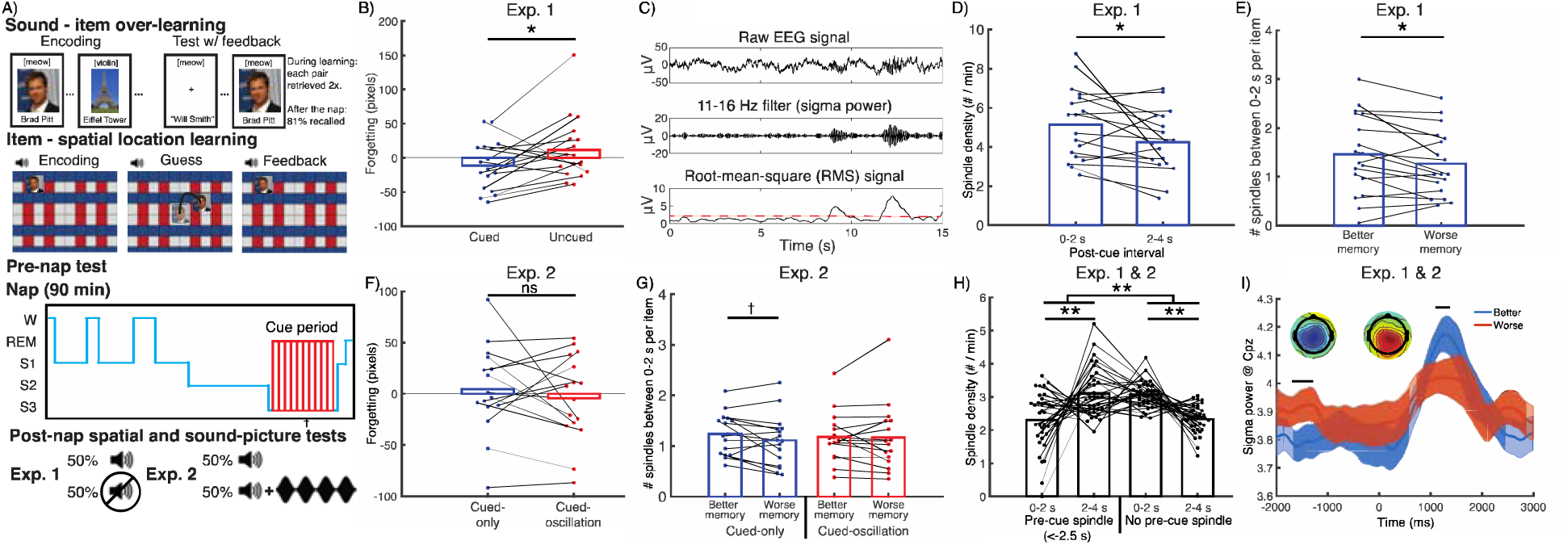
Experiments 1 & 2 procedure and results. (A) Subjects over-learned associations between sounds and items before learning item spatial locations. In experiment 1, half of the sounds were presented during slow-wave sleep (SWS) of an afternoon nap. In experiment 2, all sounds were similarly presented, and half were followed by 15-Hz, oscillating white noise (cued-oscillation). (B) In experiment 1, cues reduced forgetting between pre-nap and post-nap tests. (C) Sleep spindles calculation. (D) During the period immediately after cues (0-2 s) relative to later (2-4 s), spindles were increased. (E) A median split analysis showed more early spindles per item predicted better memory. (F) In experiment 2, forgetting did not differ between cued-only and cued-oscillation conditions. (G) Early (vs. late) spindles marginally indexed better memory for the cued-only condition, but not the cued-oscillation condition. (H) Analyses combining both experiments revealed that pre-cue spindles (occurring −2.5–0 s) reversed the prevalence of early versus late post-cue spindles. (I) Better-remembered items showed higher postcue but lower pre-cue sigma power than worse-remembered items. (Inset) Topographical maps of RMS values for better – worse memory centered around −1550 ms and 1300 ms, respectively. †: p <= 0.1. *: p <= 0.05. **: p < 0.01.

We analyzed spatial forgetting by subtracting pre-nap error from post-nap error and regressing out pre-nap error, to produce a memory change score (Fig S1). As expected, cues improved memory (*11, 12*), with less forgetting for cued than uncued items [*t*(17) = 2.2, *d_z_* = 0.5, *p* = 0.039; Fig 1B].

We next investigated relationships between post-cue spindles and memory. We detected spindles by band-passing sigma (11-16 Hz), calculating root-mean-square (RMS) values using sliding 200-ms intervals, and extracting above-threshold segments (*Methods;* Fig 1C). We found that cues tended to provoke spindles, in line with previous findings (*13, 14*). Spindles at scalp location CPz increased early relative to later after cues [0-2 s vs. 2-4 s, respectively; *t*(17) = 2.3, *d_z_* = 0.54, *p* = 0.03; Fig 1D]. Furthermore, median-split analyses revealed that more spindles occurred <2 s after better-retained items than after worse-retained items [*t*(17) = 2.23, *d_z_* = 0.53, *p* = 0.039; Fig 1E]. As in several prior studies (*15–18*), post-cue spindles positively predicted memory.

We next sought to move beyond this correlative evidence, by manipulating the incidence of spindles and seeing whether conditions that boost spindle occurrence also boost subsequent memory. First, we attempted to boost spindles directly using sensory-entrainment methods. Subjects in experiment 2 (*N*=16) heard cues during sleep, half presented alone (cued-only) and half followed by 2 s of white noise amplitude-modulated at spindle frequency (15 Hz; cued-oscillation). Given that such white-noise oscillations were previously found to facilitate spindles (*14*), we hypothesized that cued-oscillation sounds would show greater post-cue spindles as well as better retention for those cued items. Contrary to our expectations, cues with entrainment versus cues presented alone were not associated with differences in either incidence of post-cue spindles [*t*(15) = 0.20, *d_z_* = 0.05, *p* = 0.84] or corresponding memory performance [*t*(15) = 0.8, *d_z_* = 0.2, *p* = 0.44; Fig 1F]. Median-split analyses revealed a trend for more spindles for better than worse memory in the cued-only condition [*t*(15) = 1.78, *d_z_* = 0.44, *p* = 0.096], but not the cued-oscillation condition [*t*(15) = 0.13, *d_z_* = 0.03, *p* = 0.90; Fig 1G].

Since the cued-oscillation approach did not work, we next explored whether spindles were modulated by the time since the prior spindle occurrence. Prior *in vitro* (*19*), *in vivo*, (*20*), and human EEG (*21*) evidence of a decline in spindle incidence shortly after a spindle could be indicative of a refractory period. If this refractory period exists, it could be leveraged to manipulate the probability of spindle occurrence. As an initial probe into this question, we asked whether presence of a spindle pre-cue influenced post-cue spindle probability. Combining data from experiments 1 and 2, we found a significant interaction between whether there was a precue spindle (-2.5 to 0 s) and early-versus-late post-cue spindle rate [0-2 s vs. 2-4 s, respectively, *F*(1,33) = 24.23, *p* < 0.001]. Follow-up analyses connected pre-cue spindles with more late than early post-cue spindles [*t*(33) = 3.1, *d_z_* = 0.53, *p* = 0.004], and their absence with more early than late post-cue spindles [*t*(33) = 5.5, *d_z_* = 0.94, *p* < 0.001; Fig 1H]. We then asked whether pre-cue spindle activity negatively affected memory by analyzing cue-locked sigma power at CPz while correcting for multiple comparisons. As expected, sigma power was higher for better-than worse-remembered items for a post-cue time segment (1092 to 1372 ms, *p* < 0.05), but higher for worse-than better-remembered items during a pre-cue time segment (-1696 to −1288 ms, *p* < 0.05; Fig 1I). These results hinted that memories cued within spindle refractory periods may be unlikely to undergo reactivation.

To further understand these spindle refractory effects, we computed the inter-spindle lag or ISL (Fig 2A). ISL analyses showed fewer spindles for lags of 0.5-2.5 s (15.3%) compared to 2.5-4.5 s (30.0%). A parallel analysis in the frequency domain revealed a peak at 0.21 Hz (4.8-s ISL) with a range around the peak at 0.17-0.33 Hz (3-5.9-s ISL; Fig 2B). These results revealed that sigma activity occurs not randomly but rather in an oscillatory fashion.

**Figure 2.**
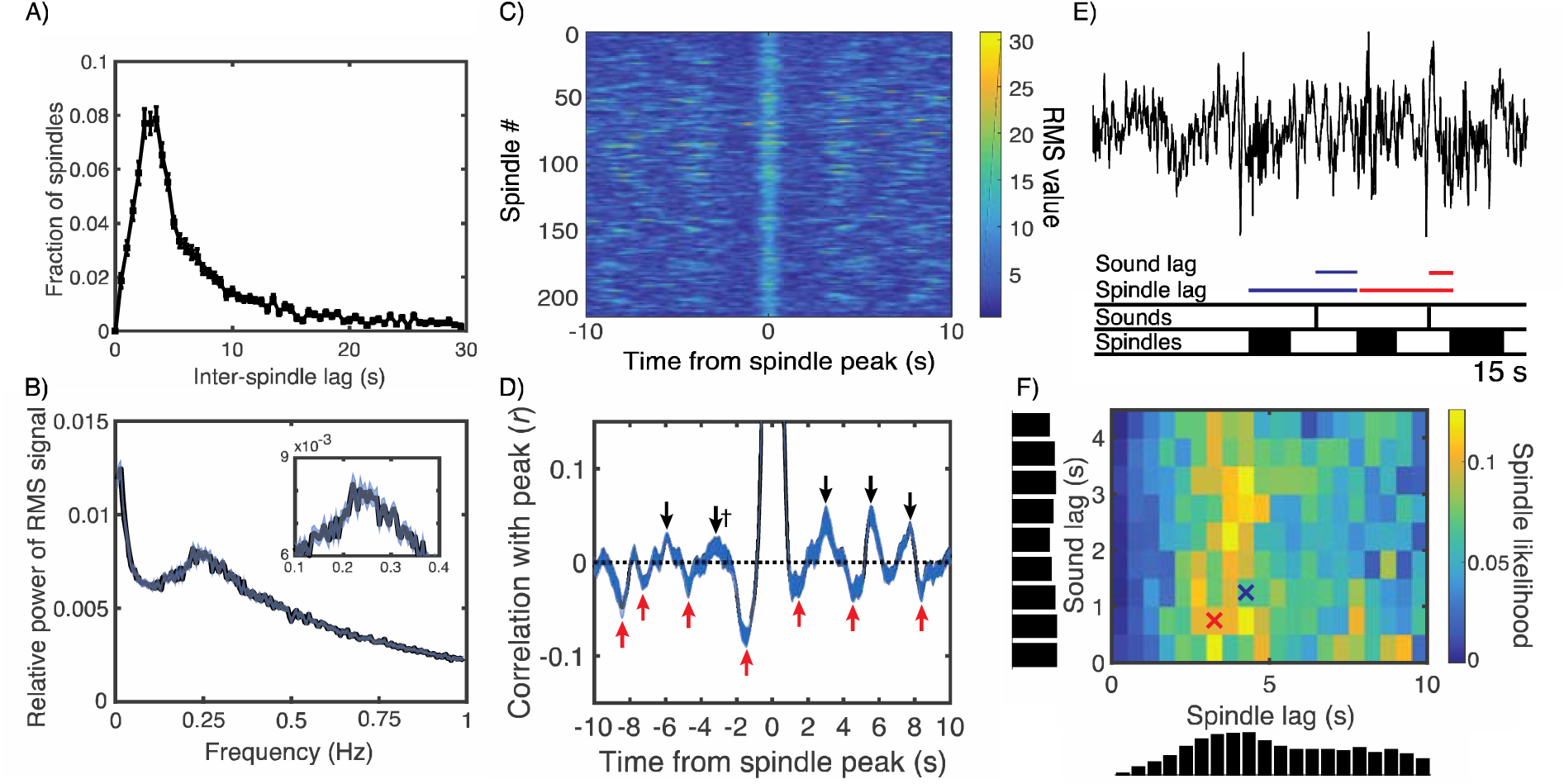
Characterizing the spindle refractory period. (A) Inter-spindle lags (ISLs) were calculated as the time between the start of successive spindles, shown for up to 30 s. (B) Fast Fourier transformations of the sigma RMS signal revealed cyclic activity most prominent in the 0.17 – 0.33 Hz range, corresponding to ISLs in the range of 3 – 5.9 s. (C) RMS values spanning from −10 s to +10 s surrounding spindle peaks from one sample subject. (D) Correlations between the RMS value of the spindle peak and all other values from −10 to 10 s were calculated for each subject. The correlation was *r* = 1.0 at the *t* = 0 spindle peak. All positive (black) and negative (red) peaks that significantly differed from zero across subjects (*p* < 0.05) are marked with arrows, except † indicates *p* = 0.07. Brackets indicate approximately symmetric peaks across *t* = 0. (E) For each instant along the recording, we found whether a spindle started or not and the time lags since the last spindle and last sound cue. (F) Likelihoods of spindles starting as a joint function of spindle and sound lag. Crosses indicate bins corresponding to color-coded time lags from (E).

Next, we obtained sigma RMS values −10 to 10 s around the RMS peaks of verified spindles (Fig 2C), and then performed autocorrelations between each time lag and the RMS peak for each subject. The autocorrelation graph revealed “reverberations,” such that on each side of *t* = 0 there were negative, positive, negative, and positive peaks (approximately ±1.5, 3.1, 4.1, and 5.35 s, respectively; Fig 2D; Table S2). To ensure these qualitative results were not artifacts of cueing, we analyzed data from subjects who did not receive cues (*N*=28; *Methods;* Fig S2). All major aspects of the above analyses held, except reverberation cycles were slightly wider (~ 4 s rather than ~ 3 s). Finally, as both preceding spindles and sounds affect spindle occurrence, we measured the likelihood of a spindle relative to a preceding sound or spindle (Fig 2E). In line with previous analyses, spindles were most likely 3-5 s after predecessors and shortly after cues; sounds shortly after spindles elicited few spindles, whereas spindles frequently occurred 3-5 s after predecessors almost regardless of cues (Fig 2F), suggesting the spindle refractory period imposes a limitation on spindle probability that can over-ride sensory influences.

If this is true, then it should be possible to manipulate spindle probability by manipulating the timing of the cue relative to the last spindle: cues presented in the refractory period should be less likely to trigger spindles and should lead to worse memory, relative to cues presented outside of the refractory period. To test this, we arranged to systematically deliver cues at different times relative to the spindle-refractory period. For experiment 3 (*N*=20), we developed an algorithm to track spindles in real-time (Fig 3A). Each cue was presented either shortly after a spindle finished (0.25 s, Early Condition) or much later (~ 2.5 s, Late Condition). Confirming the success of the spindle-tracking method, early cues showed far more pre-cue spindles than late cues [40.4% vs. 5.7%; *t*(19) = 13.3, *d_z_* = 2.97, *p* < 0.001; Fig 3B]. Crucially, spatial memory was more accurate with late cues compared to early cues [*t*(19) = 3.2, *d_z_* = 0.7, *p* = 0.004; Fig 3C]. That is, memory reactivation was apparently reduced within the spindle refractory period.

**Figure 3.**
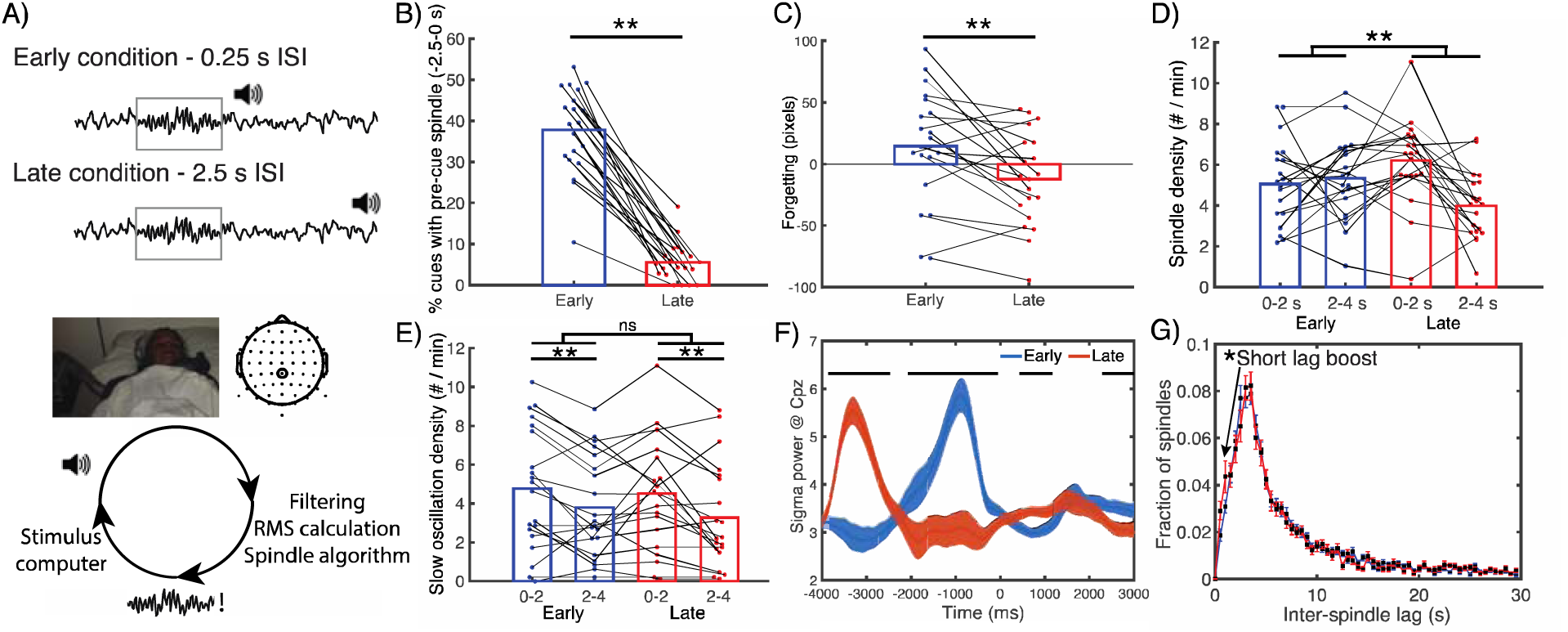
Memory retention was impaired for items cued inside versus outside the spindle refractory period. (A, top) In experiment 3, TMR cues were presented either early or late after a spindle was detected, placing them (respectively) inside or outside the spindle refractory period. (A, bottom) Real-time schematic. EEG traces from electrode CPz were filtered between 11-16 Hz, the RMS signal was calculated, and detected spindles were sent to another computer to present cues at the appropriate times. (B) A manipulation check showed a higher percentage of pre-cue spindles (<2.5 s) in the early than late cued conditions. (C) Late cues significantly enhanced memory retention relative to early cues.(D) Late cues showed a significant spindle boost early in the post-cue interval, whereas early cues did not. (E) Both early and late cues boosted slow oscillations, but there was no interaction between conditions. (F) Cue-locked analyses revealed higher early postcue sigma power for late relative to early cues. All horizontal bars indicate *p* < 0.05 segments. (G) Comparison of inter-spindle lags in experiments 1 and 2 vs. experiment 3 revealed an increased number of very short inter-spindle lags. **: *p* < 0.01.

To verify the predominance of spindles shortly after late cues, we submitted spindle-density measures to a cue type (early vs. late) x post-cue interval (0-2 vs. 2-4 s) ANOVA. We found a significant interaction between cue type and post-cue interval [*F*(1,19) = 6.6, *p* =0.01; Fig 3D]. Spindles increased early post-stimulus-onset (vs. late) after late cues [*t*(19) = 4.0, *d_z_* = 0.88, *p* < 0.001], but not after early cues [*t*(19) = 0.44, *d_z_* = 0.10, *p* = 0.66]. As stimulus-evoked slow oscillations (SOs) predict memory retention (*22*) and have also been shown to have refractory periods (*23, 24*), we next wanted to verify that effects that are attributed here to the spindle refractory period were not due instead to the SO refractory period. The same analyses on SOs, as measured using established algorithms (*25*), revealed more SOs 0-2 vs. 2-4 s after cues [*F*(1,19) = 11.66, *p* < 0.001], but no effect of our early vs. late cueing manipulation [*F*(1,19) = 2.8, *p* = 0.11] or interaction [*F*(1,19) = 0.6, *p* = 0.44; Fig 3E], indicating that the observed memory effects of this manipulation are unlikely to be due to modulation of SOs.

The reverberatory nature of sigma power was also evident in temporal patterns of cue-locked sigma RMS (Fig 3F): late cues, higher from −3868 to −2472 ms; early, higher from −1992 to −48 ms; late, crucially higher from 500 to 1172 ms; early, higher from 2252 to 3000 ms (all corrected for multiple comparisons). Finally, as *in vivo* evidence suggests the refractory period is not absolute (*20*), we reasoned that early cues directly after spindles should still occasionally able to elicit spindles, which would show up as an increase in spindles at very short ISLs. In keeping with this idea, we found higher proportions of spindles at very short lags (< 1.5 s) in experiment 3 than experiments 1 and 2 (*t*(52) = 2.38, *d* = 0.67, *p* = 0.02; Fig 3G).

Converging evidence thus describes a novel rhythm of spindle incidence. Moreover, this previously under-characterized regularity in sigma, corresponding to a ~3-5 s ISL, is strongly connected to memory reactivation. At ISL<3 s, the state of local thalamocortical spindle networks is presumably refractory due to the hyperpolarization-activated current, *I_h_* (*19, 26*). At ISL>5 s, the reduced spindle likelihood (right side of Fig 2F) may reflect the initial shift to a sleep state not conducive to spindles. Additionally, we found a strong peak at an infraslow frequency (~0.02 Hz; Fig 2B), in line with a recent report suggesting the presence of spindle-rich intervals separated by ~50 s (*27*). Our findings therefore underscore the importance of considering spindles on the meso-scale (~0.2-0.3 Hz), intermediate between the sub-second scale of the oscillations themselves (11-16 Hz) and the infra-slow scale just under a minute (0.02 Hz).

These findings parallel and qualify recent investigations into memory reactivation with respect to SOs. Given that SOs nest spindles and ripples (*7*), they could be seen to segment and thereby regulate memory reactivation. In humans, boosting SOs improves memory (*22, 28*); however, the degree to which it improves memory may be limited to effects on spindles (*23, 25*). Here we provide additional evidence that reactivation may be more broadly regulated by the spindle refractory period (~3-5 s) than individual SOs (~1s). Additionally, whereas stimulation to enhance SOs (*23, 25*) and TMR (*29*) are most effective at particular SO phases, our findings show that TMR is most effective at particular “phases” of the meso-scale sigma power rhythm of 0.2-0.3 Hz. They also suggest that efforts to manipulate spindles via optogenetic (*30*) or auditory (*14*)stimulation may also benefit by considering this rhythm.

In conclusion: Our findings suggest spindles segment sleep into prime opportunities for reactivation interspersed by gaps corresponding to the spindle refractory period. We speculate that this segmentation could serve as a mechanism for segregating memory reactivation related to different events. Furthermore, these results suggest fundamental limitations on the amount of reactivation that can occur across a given sleep period.

## Acknowledgments

This work was supported by the NIH F31-MH100958 and CV Starr fellowship to JWA, and NSF grant BCS-1461088 to KAP and KAN. We thank Neggin Keshavarzian for additional help collecting data, and Nick Depinto with help with the real-time EEG setup. Pre-registered methods for experiment 3 along with data from all experiments at the time of publication can be found at https://osf.io/brndg/. J.W.A., L.P., K.A.N., & K.A.P. conceived experiment 1. J.W.A. conceived experiments 2-3. M.W. contributed heavily to the real-time EEG code. J.W.A., M.W., & P.P. collected the data. J.W.A., L.P., K.A.N., and K.A.P. analyzed the data. J.W.A., K.A.N., & K.A.P. wrote the manuscript. All authors discussed the results and revised the paper.

## Materials and Methods

Based on the results of experiments 1 and 2, the hypotheses, methods, and planned analyses for experiment 3 were pre-registered at https://osf.io/brndg/. All code and results will be placed there upon publication.

### Subjects

Subjects were asked to wake an hour earlier than normal in order to increase the chances of their napping successfully. They were also asked to refrain from drinking alcohol the night before and caffeine the morning of the experiment. Informed consent was obtained before and monetary reimbursement given after the study. Twenty-one (*M* = 21.8 yrs, range: 18-33, 10 female) and twenty-five volunteers (*M* = 22 yrs, range: 19 - 33, 15 female) from the Northwestern community participated in experiments 1 and 2, respectively. Twenty-six volunteers (*M* = 21.2 yrs, range: 18-33, 10 female) from the Princeton University community participated in experiment 3. In experiments 1 and 2, participants’ data were included in the final dataset if they experienced one full round of cues during sleep, and excluded otherwise. In experiment 3, the rate of cue presentation was slower because we waited to detect spindles before cuing; this, in turn, made it less likely that we would be able to get through a full round of cues. Therefore, we loosened our inclusion criterion *a priori* for experiment 3: Participants were included if they experienced at least 50% of the overall cues (see Preregistered methods at https://osf.io/brndg/). Only two subjects received fewer than one full round of cues. In these cases, only the cued items from each group were analyzed. Data were excluded from subjects who did not receive the minimum number of cues (three, nine, and six subjects in experiments 1, 2, and 3, respectively).

### Stimuli

Subjects learned to associate 24 celebrities and 24 landmarks with 48 randomly-assigned environmental sounds bearing no relation to the pictures (e.g. a cat’s meow, violin musical tones). The sounds lasted 0.5 s or less and were selected from a larger set used in another TMR study (*12*), so as to maximize the distinctiveness of the selected cues from each other.

In experiment 2, the oscillating sounds were created by modulating the amplitude of a white noise signal, which was a mixture of sound frequencies from 20-1000 Hz with random amplitudes constant across the power spectrum (Antony & Paller, 2017). The modulated sound alternated between 100% and 20% of the original amplitude in the form of a sine wave using the Tremolo function in Audacity software. Thus, the modulation did not change the maximum amplitude of the signal.

### Design

The three experiments included sound-item association over-learning, item-location learning, pre-nap location testing, napping for 90 min, and post-nap item-location and sound-item testing. There was an additional phase before learning in which subjects viewed a different group of celebrities, landmarks, common objects, scrambled faces, and scrambled places, included to allow for training a wake EEG classifier. Those classification analyses are not described further in this article. The procedural details for each phase were as follows. During sound-item over-learning, subjects learned pairwise associations between 48 unique sounds and 48 new faces or places to a high degree of accuracy. During item-location learning, subjects learned the location of each picture against a background grid accompanied by its previously-associated sound. For the pre-nap test, subjects were tested on each picture location once just before sleeping, receiving feedback on the correct location after their guess. Then, subjects took a 90-min nap, during which learned sounds were randomly and repeatedly played to the subjects upon their entering SWS. Subjects returned to the lab 150 min after the nap to take tests on all item-location and sound-item associations.

### Procedure

#### Pre-training

We fitted subjects with a 64-channel cap of electrodes along with two EEG mastoid electrodes and one EMG electrode on the chin. Two EOG channels were used for monitoring horizontal and vertical eye movements.

#### Sound-item overlearning

Subjects encoded sound-item pairs using repeated study-test cycles to ensure sounds would reliably elicit picture-related neural activity. During encoding, subjects viewed each picture for 4 s; concurrent with picture onset, a unique sound was played for 0.5 s. Sound-item mappings were random and different for each subject. A label indicating the correct name and spelling of the picture was shown below 2 s after picture onset, along with a representation of the sound. During testing, subjects heard each sound alone and were asked to type in the corresponding picture label. Typing was only allowed to begin 1 s after sound offset. After submitting a response, subjects received feedback on whether their response was correct, along with a 4-s presentation of the picture and label, with the sound presented twice at 0 and 2 s. After the subject correctly produced a label twice in a row, the corresponding pair was dropped from further testing. Testing continued until all labels were correctly produced twice in a row. The pace of encoding and testing were thus at the subject’s discretion in this phase.

#### Item-location learning

Next, subjects learned the location of each item against a background grid. Each picture was randomly assigned to a location −300 to 300 pixels from the center of the screen in horizontal and vertical directions. During encoding, subjects viewed the location of each picture for 3 s, accompanied by a single presentation of the picture’s accompanying sound. Following encoding, we asked subjects to drag each picture from the center of the screen to where they remembered seeing it. After subjects made their location response, they viewed feedback of each picture in its correct location and heard its corresponding sound. When the guessed location occurred within 150 pixels of the correct location, the item dropped out from further testing. Subjects performed the task until they guessed each pair within 150 pixels once.

#### Pre-nap item-location test

Following a 5-min break, subjects made their location response for each item once, followed by feedback. Items were again accompanied by their corresponding sound at presentation and feedback.

#### Nap

Subjects then took a nap in the laboratory against a background white noise level of approximately 40 dB. Following online indications of SWS, 0.5-s sound cues were administered once every 4.5 s in a randomized order. In experiment 1, half of the cues were presented over multiple rounds (*M* = 7.24, range = 2.9-9.25). In experiment 2, all sounds were presented (*M* = 6.53, range = 2.62-10.77), with half of the sounds followed by 2 s of 15-Hz oscillating white noise. In experiment 3, all sounds were presented (*M* = 1.96, range = 0.56-4.0), with half 0.25 s after the end of a spindle and half 2.5 s after the end of a spindle. In all experiments, items in each group were equally split between celebrity and landmark categories. No individual cue caused an amplitude jump greater than 4 dB. Cues were immediately stopped upon arousals and repeated if the subject remained in SWS after all cues finished. If a subject had not heard a sufficient number of cues after 60 min, we cued during stage-2 NREM sleep.

Following the nap, subjects returned after 150 min and were tested on each item-location and then each sound-item pair without feedback. Sounds were omitted from the post-nap item-location test to prevent them from influencing memory for the later post-nap sound-item pair test. At the completion of the experiment, subjects were paid for their time and debriefed about the aims of the experiment.

### Dependent variables

We used an adjusted forgetting score as our primary dependent variable. Forgetting, calculated as post-nap error minus pre-nap error, significantly correlates with pre-nap error. Items with highly accurate pre-nap recall face ceiling effects (e.g. an error of only 2 pixels cannot be improved across the nap by more than 2 pixels) and those with poor pre-nap accuracy show a regression to the mean (e.g., an incorrectly recalled location, when very distant from the correct location, is likely to be recalled more accurately after the nap, even by chance). Therefore, we calculated the linear relationship between pre-nap score and forgetting (post-nap – pre-nap score) pooled across subjects in the present data (Fig. S1). Then we subtracted each forgetting score from the forgetting expected from this linear relationship (i.e., the residual) to produce the adjusted forgetting store used for all reported analyses.

### EEG recording and pre-processing

Continuous EEG was recorded during the nap using Ag/AgCl active electrodes (Biosemi ActiveTwo, Amsterdam) using the same electrode layout and recording hardware at Northwestern (experiments 1 and 2) and Princeton (experiment 3). In experiment 3, for the purposes of real-time analyses, EEG data were collected using OpenViBE rather than Biosemi software. Recordings were made at 512 Hz from 64 scalp EEG electrode locations. In addition, a vertical electrooculogram (EOG) electrode was placed next to the right eye, a horizontal EOG electrode was placed under the left eye, and an electromyogram (EMG) electrode was placed on the chin.

EEG data were processed using a combination of internal functions in EEGLAB (Delorme & Makeig, 2004) and custom-written scripts. Data were re-referenced offline to the average signal of the left and right mastoid channels and were down-sampled to 256 Hz. They were high-pass filtered at 0.1 Hz and low-pass filtered at 60 Hz in successive steps. Problematic channels were interpolated using the spherical method.

### Sleep physiological analyses

Sleep stages were determined by an expert scorer according to standard criteria (*32*) Table S1 shows the breakdown of stages for each condition as well as the number of cues occurring within each stage for all experiments. Note that sleep-staging rules require assigning stages based on whichever stage is more prevalent within the 30-s epoch, which can result in sounds occurring in stages that were not the intended targets. Artifacts (large movements, blinks, arousals, and rare, large deflections in single channels) during sleep were marked separately in 5-s chunks following sleep staging.

Spindles and slow oscillations were calculated using established algorithms. Each of these scripts ignored 5-s intervals marked for rejection, effectively stitching together non-artefactual segments from all NREM epochs into a long, continuous EEG signal for further processing.

For spindles, sleep EEG data were bandpass-filtered between 11-16 Hz using a two-way, least-squares finite-impulse-response filter. Next, we calculated a root-mean-square (RMS) value for every time point using a moving window of ±0.2 s for each channel separately. A threshold was determined by multiplying the standard deviation of the entire channel’s signal by 1.5 (*33*). Any RMS signal that crossed this threshold consecutively for 0.5- to 3-s was considered a spindle. Times for the start, negative peak (largest negative voltage value), and end of each spindle were recorded for alignment with sleep cues. We used this same RMS calculation for online spindle detection and offline cue-locked analyses.

For counting slow oscillations, sleep EEG data were first low-pass filtered at 3.5 Hz. Any series of data points with successive positive-to-negative crossings lasting 0.75 to 2 s (corresponding to 0.5-1.3 Hz), a negative peak of −40 μV, and peak-to-peak amplitude of 75 μV was considered a slow oscillation. Similar to spindles, the start, negative peak, and end were recorded for later alignment with sleep cues.

As fast spindles tend to correlate with subsequent memory (*2*), we chose a cluster of centroparietal electrodes (Cz, Cp1, Cpz, Cp2, Pz) for physiological analyses *a priori* as they are the scalp locations where fast spindle power is maximum (*34*). Cue-locked spindle density measures (e.g. 0-2 s, 2-4 s) indicate spindles started for all relevant periods divided by the length of these periods in minutes. We chose electrode CPz where single channels were more appropriate, such as graphing RMS over time, as well as for our online spindle detection algorithm. Fig. 1I uses non-baseline-corrected RMS values to show pre-cue effects; Fig. S3 depicts the same analysis using baseline correction (by subtracting the mean of the interval between −0.5 to 0 s). We corrected for multiple comparisons in two steps. First, we randomly permuted better-remembered and worse-remembered conditions 400 separate times. After each permutation, we calculated the maximum number of consecutive time values that differed from *p* < 0.05, yielding a null distribution of maximum-consecutive-timepoints. We then compared the number of consecutive significant timepoints to the null distribution, yielding a family-wise error value. Any true time segment exceeding the 95^th^ percentile of the null distribution (indicating a family-wise *p* value < 0.05) was deemed significant.

### Spindle refractory period analyses

All analyses in Fig. 2 were performed on electrode CPz. Inter-spindle lags were found by calculating the amount of time between successive spindles. Relative RMS power was calculated by performing fast Fourier transforms on artifact-free NREM periods, sorting data into 200 frequency bins between 0-1 Hz, and calculating within-subject relative power. To calculate RMS autocorrelations, we first obtained all RMS values between −10 to +10 s surrounding each spindle peak for each subject, mean-normed for the average RMS values from a wider −30 to +30 s block. Next, we calculated correlations between every time lag and *t* = 0 separately across spindle trials. Finally, we plotted these mean correlations with standard errors with arrows where correlations were both at a local maxima or minima and significantly different from zero across subjects. Bivariate spindle and sound lag analyses were calculated by taking each moment in the recording and binning it into 0.5-s segments by the amount of time since the onset of the most recent sound and the onset of the most recent spindle. Moments when spindles started were marked to later calculate the likelihood of a spindle occurring in that given bivariate bin. Finally, two-dimensional color plots were produced for sound intervals of 0 – 4.5 s (9 bins) and spindle lags of 0 – 10 s (20 bins), and horizontal and vertical bar graphs indicated the means across each dimension.

### Real-time spindle algorithm

In order to time cues relative to spindle events, we created an online spindle detection algorithm using open-source brain computer interface software called OpenViBE (http://openvibe.inria.fr/). OpenViBE allows for real-time processing of EEG data through MATLAB scripting, and the translation of established offline detection scripts to an online format. Our offline algorithm used a band-pass (11-16 Hz) filter, RMS values based on a ± 0.2-s moving window, and a single-threshold constant value to detect spindles based on the standard deviation of the RMS signal over the whole recording.

However, in real-time, one cannot use the whole recording to calculate baseline thresholds or reliably reject artifacts. Therefore, we used a different algorithm for our online scripts (Fig. S4). We relied on two band-pass filters and two spindle thresholds, instead of one for each. For filters, we chose the sigma band (11-16 Hz) and an equal-sized lower beta band that should have no spindle information (16-21 Hz). Generally, spindles only occur in the sigma band, but artifacts show up in both bands. As such, monitoring lower beta allowed us to detect (and discount) periods of time when broader signal artifacts were present. This algorithm was chosen based on its performance in detecting spindles on an online sleep spindles database (http://www.tcts.fpms.ac.be/~devuyst/Databases/DatabaseSpindles/).

The lower and upper thresholds were 2 and 4.5 times the RMS mean of the last 600 s of the recording for the baseline (lower beta) frequency bandwidth. If the RMS values in the spindle frequency bandwidth crossed the lower threshold, it was a candidate for a spindle. The length of time the RMS signal was above this lower threshold constituted its duration. To be categorized as a spindle, the duration was required to be 0.5-3 s and the RMS value was required to surpass the upper threshold at least once. Thus, if the spindle RMS values crossed the first minimum threshold without reaching the second maximum threshold, then it would not be categorized as a spindle, no matter its duration. Similarly, if it crossed the maximum threshold but did not have a duration between 0.5-3.0 s, it would also not be categorized as a spindle.

In order to present sounds, three conditions needed to be met: (1) spindle detection, (2) RMS values were currently below the spindle threshold (so there was no candidate spindle presently ongoing) and (3) > 4.5 s elapsed since the onset of the previous sound. Once these conditions were met, sound information was sent to the presentation computer via UDP packets at different times depending on sound type. Early intervals of 0.25 s were chosen (as opposed to no delay) in order to avoid a scenario in which RMS values, while below the spindle threshold, still remained above baseline RMS values, thus causing a possible confound of sigma power at the *t* = 0. Late intervals of 3.5 s were chosen to be late enough to fall outside of the refractory period (see Fig. 2). If a spindle occurred during this interval, the timer was reset and the next late sound could only occur 3.5 s after the onset of that (intervening) spindle.

### Spontaneous (non-TMR) sleep data

Data collected during similar experiments from other subjects (*N*=28, age range: 18-30) were used to validate the dynamics of the spindle refractory period in the absence of TMR cues. The no-sounds control condition in a published study (*35*) was used for 16 subjects. In all cases, subjects performed pre-learning memory tasks. These data were acquired using Neuroscan software at a sampling rate of 1000 Hz with a bandpass of 0.1–100 Hz. Tin electrodes in an elastic cap were placed at 21 standard scalp locations, left and right mastoids, lateral to the right eye, under the left eye, and on the chin. Data were downsampled to 250 Hz, re-referenced to average mastoids, and filtered between 0.4-60 Hz in successive steps using a 2-way least-squares finite impulse response filter. Because we used an electrode montage that did not include CPz, we used Pz instead. Analyses paralleling Fig. 2A, B, and D are reproduced using this dataset in Fig. S2.

### Supplementary Text

#### Baseline-corrected sigma RMS from experiments 1 and 2

To highlight the role of pre-cue sigma power, we did not perform baseline correction in Fig. 1I. Nevertheless, baseline-corrected sigma power showed a similar post-cue subsequent memory difference (1052 to 1400 ms, *p* < 0.05), but predictably no pre-cue difference (Fig. S3).

#### Final sound-item memory

We tested overlearned sound-item memory after the final spatial item-location test. Even though we expected ceiling-level performance, we analyzed final sound-item memory because it is conceivable that cueing could also strengthen these associations. Overall, these results showed non-significant TMR benefits that were weaker than, but generally consistent with, the spatial error results in each experiment. In experiment 1, cued associations were marginally better-remembered than uncued associations [*t*(17) = 1.7, *d_z_* = 0.4, *p* = 0.106]. In experiment 2, no difference was found between cued-only than cued-oscillation associations [*t* (15) = 0.93, *d_z_* = 0.23, *p* = 0.37]. In experiment 3, late cued associations were marginally better-remembered than early cued associations [*t* (19) = 1.8, *d_z_* = 0.4, *p* = 0.089].

**Fig. S1.**
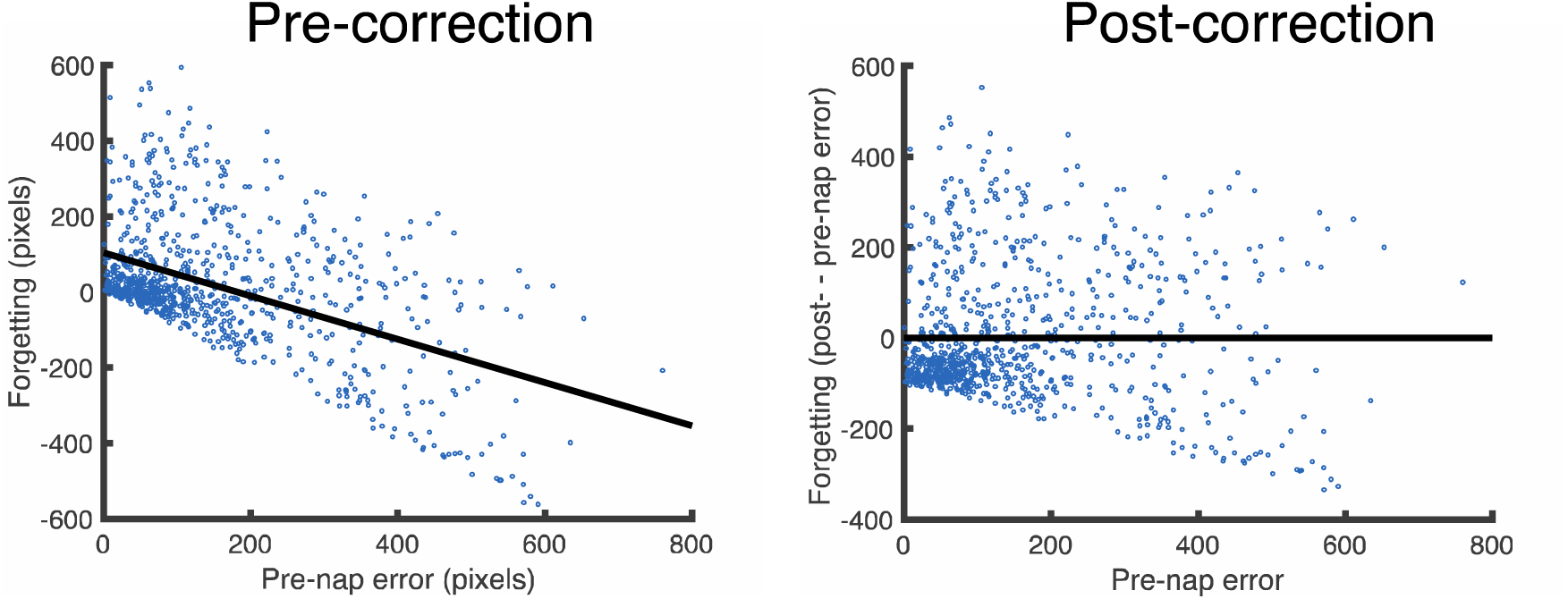
Residual analysis used in the forgetting metric. (A) Pre-nap error significantly predicts forgetting (post-– pre-nap error). (B) Corrected (residual) forgetting values after regressing out pre-nap error.

**Fig. S2.**
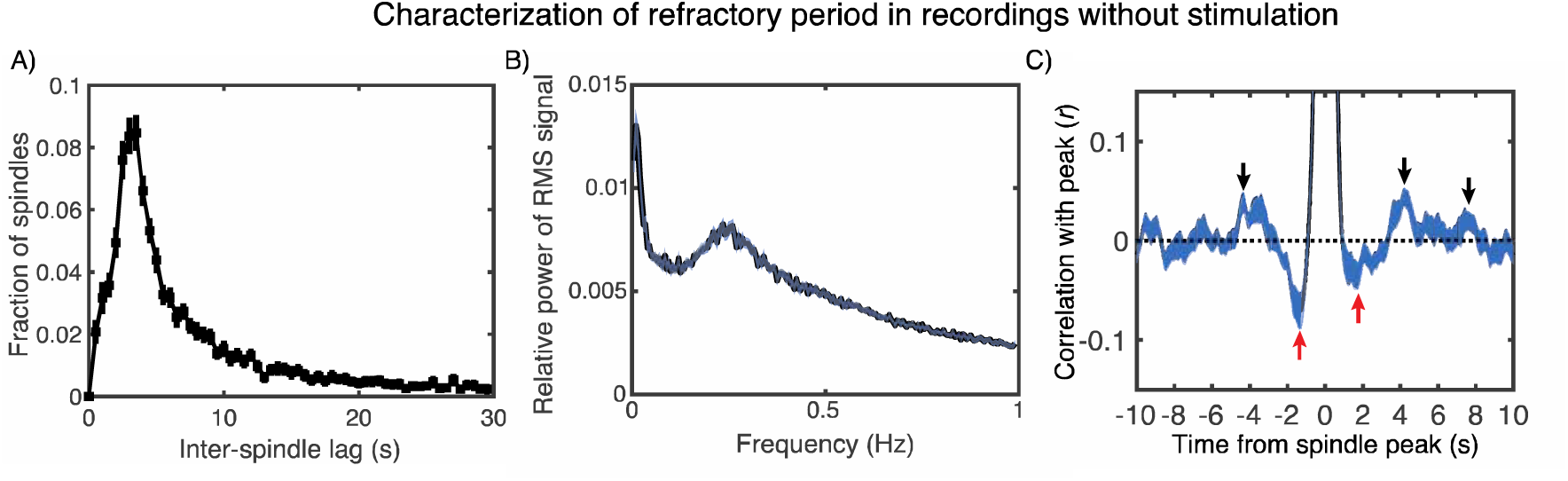
Spindle refractory period analyses on non-TMR data. We created a variant of the graphs from Fig. 2A, B, and D using data that had no TMR cues (*N*=28). We relied on data from the Pz electrode because EEG acquisition did not include CPz.

**Fig. S3.**
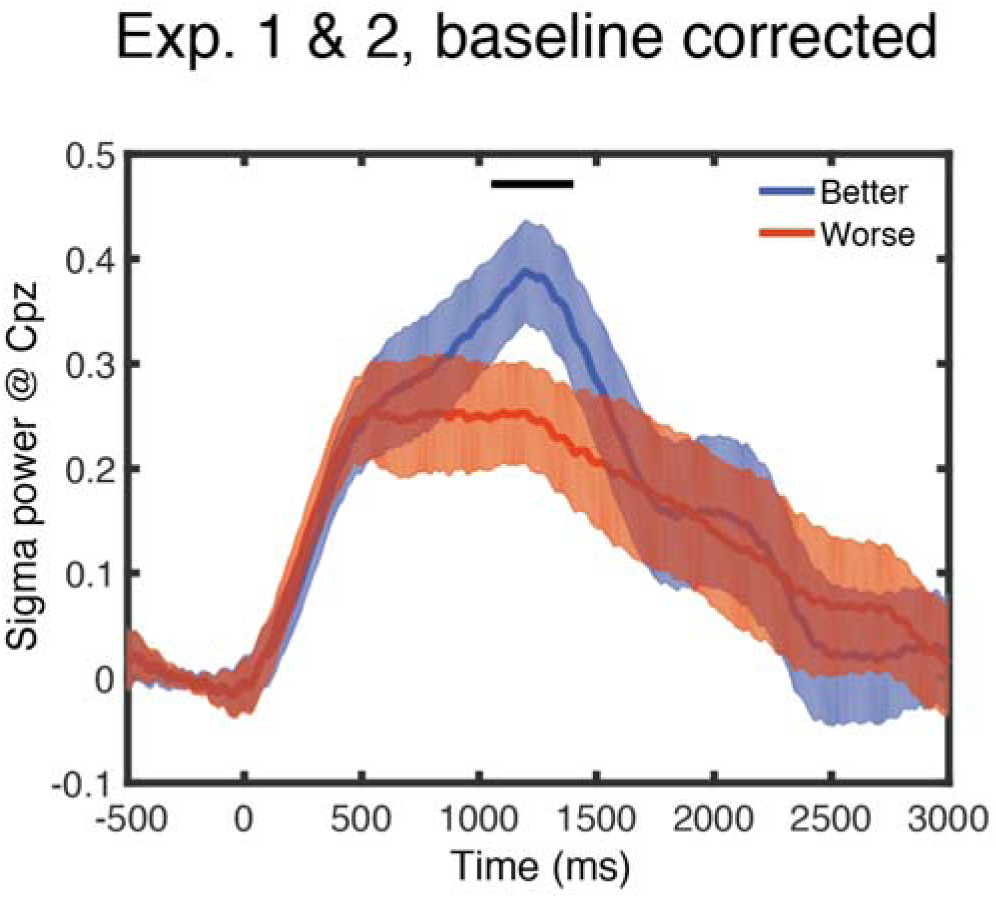
Baseline-corrected subsequent memory analyses for experiments 1 & 2. We plotted cue-locked RMS sigma values for better- and worse-remembered items, as determined by median split. The horizontal bar indicates times showing a significant difference between the conditions at *p* < 0.05, which was significant after correcting for multiple comparisons.

**Fig. S4.**
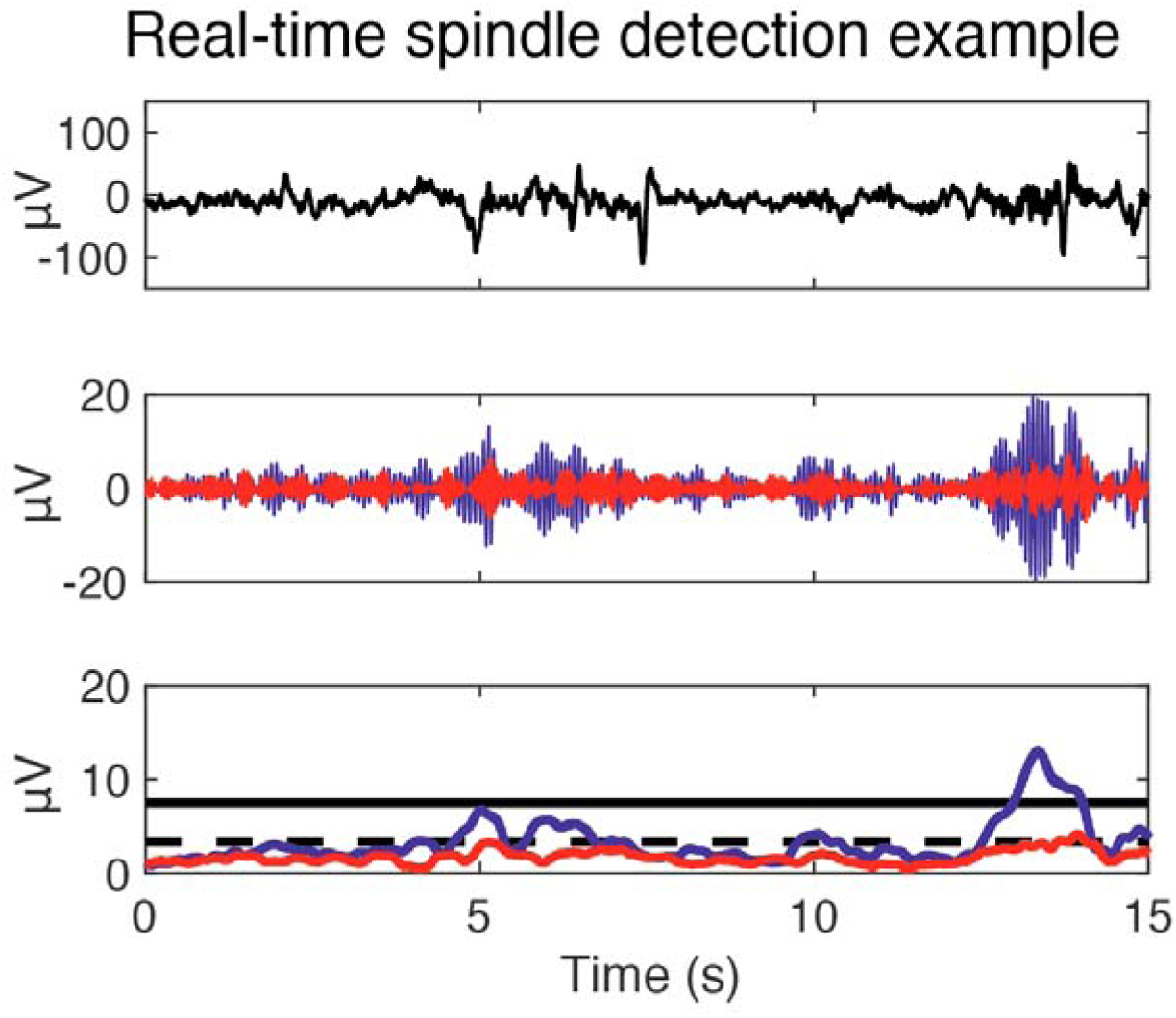
Example of a spindle detected by online spindle algorithm. (Top) Raw EEG signal. (Middle) Signals filtered in the sigma (11-16 Hz, blue) and lower beta (16-21 Hz, red) bands. (Bottom) RMS sigma (blue) and lower beta (red) signals along with the first (dashed black) and second (solid black) spindle thresholds, which were multiplications of 2 and 4.5 times the mean lower beta power, respectively. Any sigma signal above the first threshold between 0.5 – 3 s and above the second threshold at any point was considered a spindle. One spindle was thus detected near the end of the interval.

**Table S1.** Sleep staging and cue quantification. Mean amount of time in each sleep stage (min ± SEM) is displayed together with the number of cues per stage for Experiments 1, 2, and 3.

| Time in each stage (min) |  | Wake | S1 | S2 | S3 | REM |
| --- | --- | --- | --- | --- | --- | --- |
| Exp 1 | Mean | 27.69 | 5.78 | 24.25 | 20.81 | 3.08 |
|  | SEM | 3.96 | 0.69 | 2.90 | 4.20 | 1.07 |
| Exp 2 | Mean | 21.22 | 5.97 | 22.69 | 20.16 | 4.28 |
|  | SEM | 3.51 | 0.95 | 2.54 | 3.23 | 1.35 |
| Exp 3 | Mean | 21.43 | 5.83 | 26.68 | 17.45 | 4.98 |
|  | SEM | 3.59 | 0.56 | 2.49 | 2.41 | 1.53 |
| Mean sounds per stage |  |  |  |  |  |  |
| Exp 1 | Mean | 4.17 | 2.61 | 43.11 | 120.50 | 0.00 |
|  | SEM | 1.50 | 0.97 | 13.24 | 19.91 | 0.00 |
| Exp 2 | Mean | 1.81 | 1.44 | 97.31 | 217.25 | 0.00 |
|  | SEM | 0.70 | 0.83 | 26.90 | 30.97 | 0.00 |
| Exp 3 | Mean | 1.30 | 0.80 | 35.90 | 54.30 | 0.85 |
|  | SEM | 0.42 | 0.30 | 3.97 | 9.88 | 0.43 |

**Table S2.**
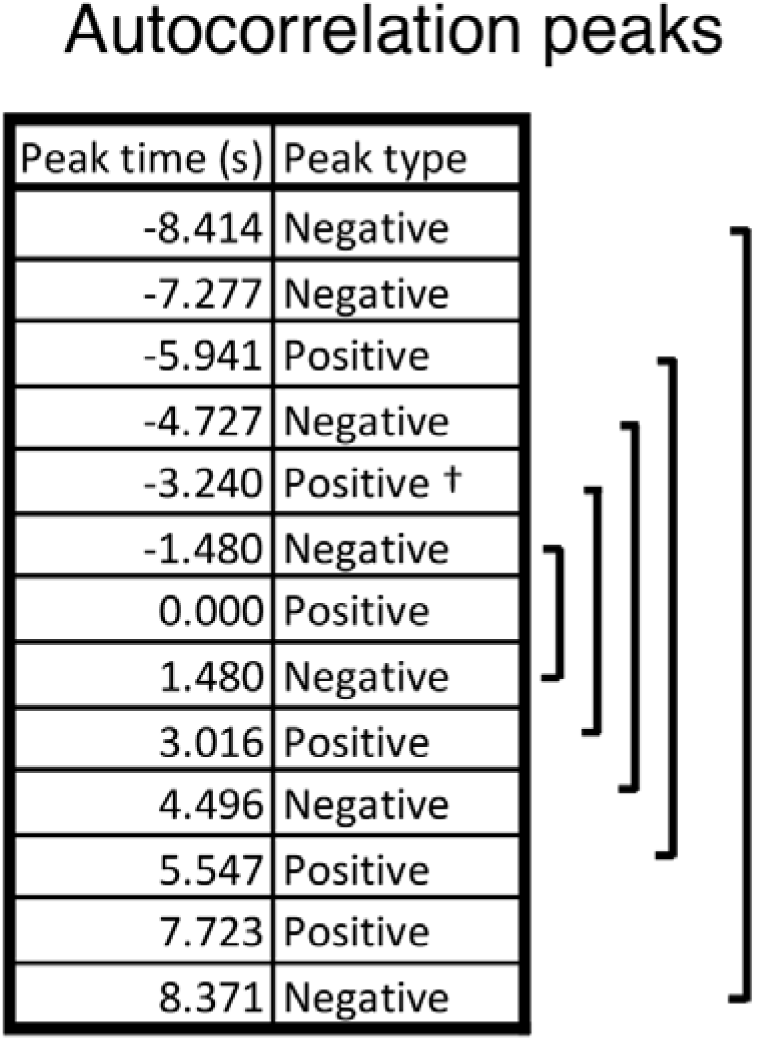
Timing of autocorrelation peaks in sigma RMS signal. Shown are all positive and negative autocorrelation peaks with each time point and *t* = 0 from Fig. 2D. Brackets indicate approximately symmetric peaks around *t* = 0. All peaks are significant at *p* < 0.05 level, except †, where 0.05 *p* < 0.1.

Autocorrelation peaks
| Peak time (s) | Peak type |
| --- | --- |
| -8.414 | Negative |
| -7.277 | Negative |
| -5.941 | Positive |
| -4.727 | Negative |
| -3.240 | Positive † |
| -1.480 | Negative |
| 0.000 | Positive |
| 1.480 | Negative |
| 3.016 | Positive |
| 4.496 | Negative |
| 5.547 | Positive |
| 7.723 | Positive |
| 8.371 | Negative |

